# HIPI: Spatially Resolved Multiplexed Protein Expression Inferred from H&E WSIs

**DOI:** 10.1101/2024.03.26.586744

**Authors:** Ron Zeira, Leon Anavy, Zohar Yakhini, Ehud Rivlin, Daniel Freedman

## Abstract

Solid tumors are characterized by complex interactions between the tumor, the immune system and the microenvironment. These interactions and intra-tumor variations have both diagnostic and prognostic significance and implications. However, quantifying the underlying processes in patient samples requires expensive and complicated molecular experiments. In contrast, H&E staining is typically performed as part of the routine standard process, and is very cheap. Here we present HIPI (H&E Image Interpretation and Protein Expression Inference) for predicting cell marker expression from tumor H&E images. We process paired H&E and CyCIF images taken from serial sections of colorectal cancers to train our model. We show that our model accurately predicts the spatial distribution of several important cell markers, on both held-out tumor regions as well as new tumor samples taken from different patients. Moreover, using only the tissue image morphology, HIPI is able to colocalize the interactions between different cell types, further demonstrating its potential clinical significance.

## 1 Introduction

Analysis of histopathological images from stained tissue sections has played a key role in identifying tumor features with diagnostic and prognostic significance [1]. Stained tumor inspection enables the characterization of intra-tumor regions and of tumor interactions with the microenvironment and the immune system. Moreover, it facilitates the detection of predictive biomarkers for disease progression and treatment [2]. Hematoxylin and eosin (*H&E*) staining of tumor tissues is routinely performed as part of cancer standard care [3]. Using H&E stained Whole Slide Images (*WSI*s) of a tumor, pathologists can distinguish between the nuclear and cytoplasmic parts of cells, and identify patterns in tissue structure and in the cell distribution. H&E staining is often complemented by immunohistochemistry (*IHC*) staining of additional tissue slices which allows the identification and quantification of cancer and immune-specific biomarkers [4]. However, IHC staining is a more delicate, time consuming and expensive process; and therefore, the number of IHC stains performed on each sample is limited. Furthermore, not all cellular molecular biomarkers can be detected using IHC staining.

Spatial molecular methods have been emerging in recent years as a powerful tool for measuring cellular biomarkers while retaining spatial information and highlighting spatial relationships [5]. Such approaches quantify the expression of different molecular modalities together with their spatial locations, providing a rich molecular image of the tissue that may not always be obtained with traditional staining methods. Data acquired using spatial technologies varies in terms of the type of modality (RNAs, proteins, metabolites), spatial resolution (sub-cellular, cellular, a few cells) and the number of measured features (a few targeted features, genome/transcriptome-wide measurements) [6]. For instance, *Spatial Transcriptomics* measures the RNA expression of all genes across thousands of spots on a tissue slice, each containing around a dozen cells [7]. In a different approach, the cyclic immunofluorescence (*CyCIF*) technology creates multiplexed images of targeted protein expression and enables the detection of cells and cellular expression [8]. Nevertheless, spatial technologies have mainly been used for exploratory cancer research, while their application for routine care is very limited due to the specialized equipment and cost required [5].

Computational pathology methods for the analysis of histopathology images have resulted in significant interest and progress recently [9, 10]. Artificial intelligence models have been trained to facilitate various tasks that previously required manual labor and expert annotation, as well as novel tasks that were not manually possible. For example, models have been developed to predict prognostic and diagnostic labels from sample WSIs such as cancer staging, grading, classification, sub-typing, survival prediction, treatment response, biomarker detection, etc. [11, 12]. Another class of models, referred to as *virtual staining*, are trained to transform one type of stained tissue WSI into a different type of stain of the same tissue, reducing the need to actually stain multiple tissues [13, 14]. More recent models allow for the transformation of histopathology images into spatial molecular maps, such as RNA expression measured using spatial transcriptomics [15, 16, 17, 18, 19]. However, training these models is difficult, as datasets of paired images and spatial molecular data from the same tissue are scarce. Therefore, these models were evaluated only on slices coming from the same tissue used for training but were mostly unsuccessful generalizing to new tissue samples [15, 16, 17]. A very recent article proposed a virtual staining model to map H&E into virtual CyCIF stains [20], but only evaluated on slides coming from the same tissue.

Here we propose an artificial intelligence model called **H&**E Image **I**nterpretation and **P**rotein Expression **I**nference (**HIPI**) to predict multiplexed CyCIF protein expression levels from whole slide H&E images. We align H&E and CyCIF data taken from adjacent tissue slices of colorectal cancer in order to train our model and predict multiple cellular markers. To cope with data scarcity, we base our model on recent advancements in Self-Supervised Learning (*SSL*) that train on large amounts of unlabeled data to extract meaningful image representations [21, 22, 23]. We further use an augmentation scheme during training to address staining intensities and other artifacts specific for H&E images.

We show that HIPI accurately predicts protein expression levels on both held-out colorectal cancer tissue regions and on new tumor samples taken from different patients not used during training. Our method is able to detect the occurrence and co-occurrence of important molecular markers based only on image morphology from routinely taken H&E WSIs. For instance, the interaction between PD1 and PDL1 markers predicted by our model can be therapeutically targeted in colorectal cancer patients and thus is clinically significant [24]. Moreover, our model enables the generation, or inference, of spatial maps for multiple markers. This inference of vector valued markers would be too expensive to generate with traditional IHC stains.

## 2 Results

### 2.1 Predicting CyCIF Protein Expression Levels from H&E Images

We developed **H&**E Image **I**nterpretation and **P**rotein Expression **I**nference (**HIPI**), a deep learning model for predicting CyCIF protein expression levels from whole slide H&E images (Figure 1a, Supplementary methods S1.2). HIPI predicts the expression levels of multiple proteins from a single H&E image, operating on image tiles. Each tile is fed into a feature extraction model pretrained on many histological images with self supervision [23]. The resulting feature vector is used as the input for a -layer fully connected regression that outputs the expression values of various proteins. The protein expression of each tile is then reordered (de-tiled) to give tissue-wide protein expression maps.

**Figure 1:**
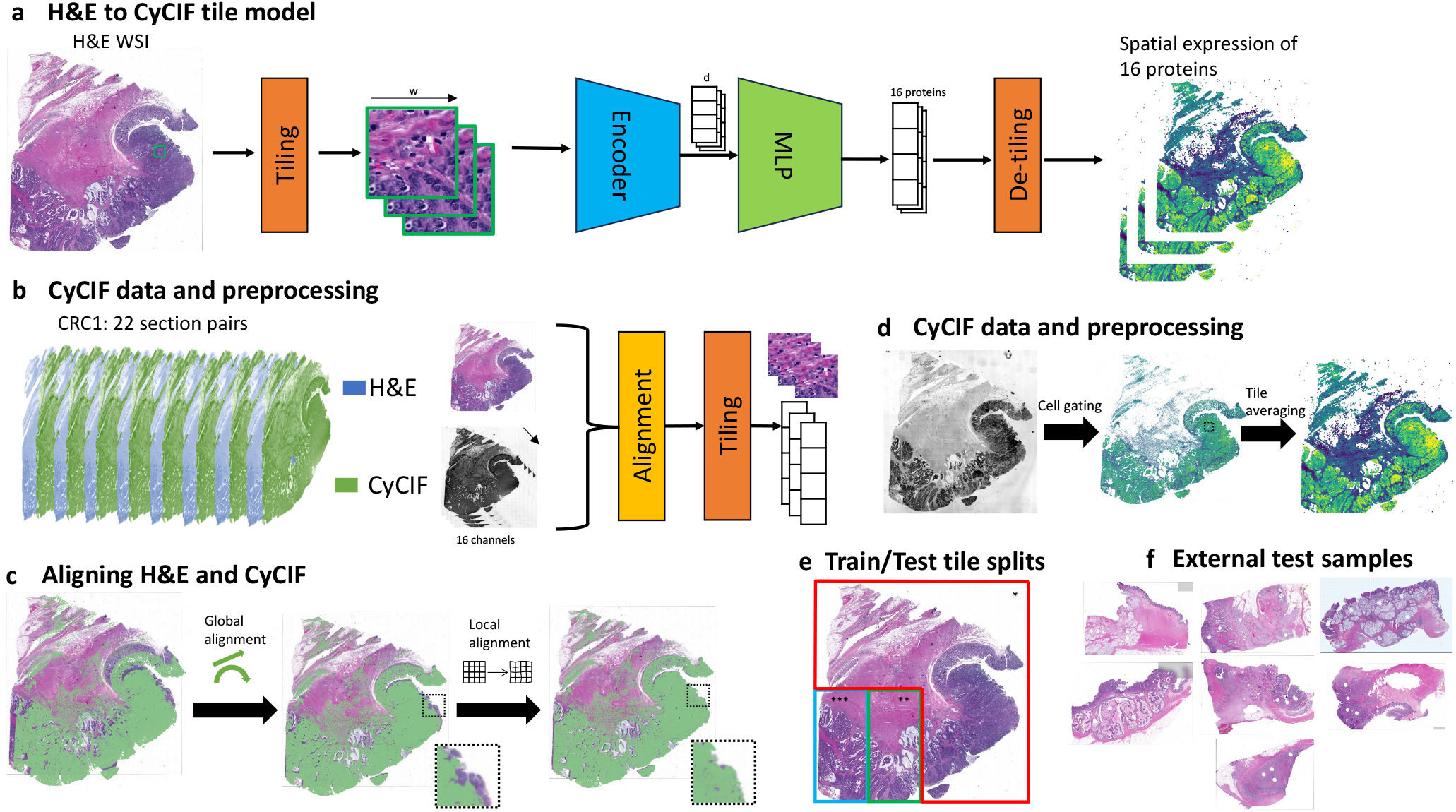
**H&**E Image **I**nterpretation and **P**rotein Expression **I**nference - overview of the data and method: (a) A schematic overivew of HIPI. An H&E slide is processed in tiles through a Deep-Learning prediction model generating multiplexed spatial expression level of 16 proteins. (b) An overview of the data and preprocessing steps. Pairs of H&E and CyCIF images taken from adjacent tissue slices are aligned and processed to generate image tiles with corresponding expression levels. (c) Alignment of adjacent H&E and CyCIF images using global linear transformation followed by local non-linear registration. The inset is a zoom-in on a tile to demonstrate the effect of the local refined alignment. (d) Deriving tile level expression from CyCIF cell data. Cell locations and expression were taken from [25]. The expression is then aggregated at the tile level. The figure illustrates expression of Ki67 marker on slice 25 (yellow - high expression). (e) Train (* red), validation (** green) and test (*** blue) data splits for sample CRC1 illustrated on a single slice. (f) Seven additional CRC samples from different patients.

We trained HIPI on a subset of tiles from 22 pairs of CyCIF and H&E Whole Slide Images (WSIs) taken from adjacent serial slices of an adenocarcinoma patient obtained from the literature (Figure 1b) [25]. This is the sample referred to as CRC01 in [25] as well as in this work. We used 16 measurements that were used by [25] as cellular markers: cytokeratin, Ki-67, CD3, CD20, CD45RO, CD4 CD8a, CD68 CD163, FOXP3, PD1, PDL1, CD31, *α*-SMA, desmin, and CD45. The training of HIPI required matching images of H&E with images of the aforementioned CyCIF protein levels. Direct mapping of cells between adjacent tissue slices is challenging due to both biological and technical variation. Therefore, we used a two-step registration process of adjacent CyCIF and H&E image pairs. First we performed a global alignment of the entire CyCIF image onto the H&E image using an affine transformation. Second, the alignment was fine-tuned using non-linear tile level local alignment (Figure 1c, Supplementary methods S1.1.1). The model was then trained on tile-level protein expression for tiles of size 256 *×* 256 pixels, each corresponding to roughly a 128*µm ×* 128*µm* tissue tile. The tile level protein expression is calculated by aggregating all corresponding cell calls from the CyCIF data (Figure 1d, Supplementary methods S1.1.2).

We trained the model using 75% of the tiles coming from the same three quadrants of the 22 tissue WSIs while the tiles in the bottom left quadrant of each slide were left out as test data (Figure 1e). Out of the bottom left quadrant tiles, the right half was used for validation and assessment of the model during training while the left half was left out for assessment of the final model as test data. With this partition, we make sure that tiles used for training and testing come from different areas of the tissue, avoiding memorization and eliminating spatial effects on the results. This is in contrast to previous models that either evaluated on adjacent slices from the same tissue or randomly split regions of the same tissue into training and test sets [15, 16, 17, 20]. Overall, the training set had 1,351,680 tiles, the validation set had 388,783 tiles and the test set had 379,142 tiles. During training we used an image augmentation scheme designed specifically for pathology images better generalization [26].

### 2.2 HIPI Predicts CyCIF from H&E

We trained HIPI to predict CyCIF protein expression from H&E image tiles from CRC01 and used the model to predict protein markers on all image tiles (Section 2.1). Overall, we observe that our model’s predictions are highly correlated to the measured expression with median Pearson correlation of 0.66, 0.61, 0.64 for the training, validation and test tiles, respectively, across all slice pairs and proteins (Figures 2 and S1). Moreover, on most markers we do not see significant differences in prediction performance between tiles used for training and the tiles left out for testing. Between the different slice pairs, we see similar levels of prediction correlation for a given protein. For some markers, e.g. Keratin, Ki67 or CD45, the model shows high concordance with the measurements with median correlations of 0.86, 0.73 and 0.67 respectively between the tiles of the different slice pairs (Figure 2). Furthermore, the model is especially successful at capturing regions with high marker expression. To show this, we calculated the *top-X% accuracy* defined as the overlap percentage between the highest X% of tiles measured for a certain marker and the highest X% of tiles predicted for the same marker. For instance, the top-20% accuracy obtained by the model in slice 96 for Keratin, Ki67 and CD45 is 72%, 60% and 65% respectively (Figure S4, Table S1).

**Figure 2:**
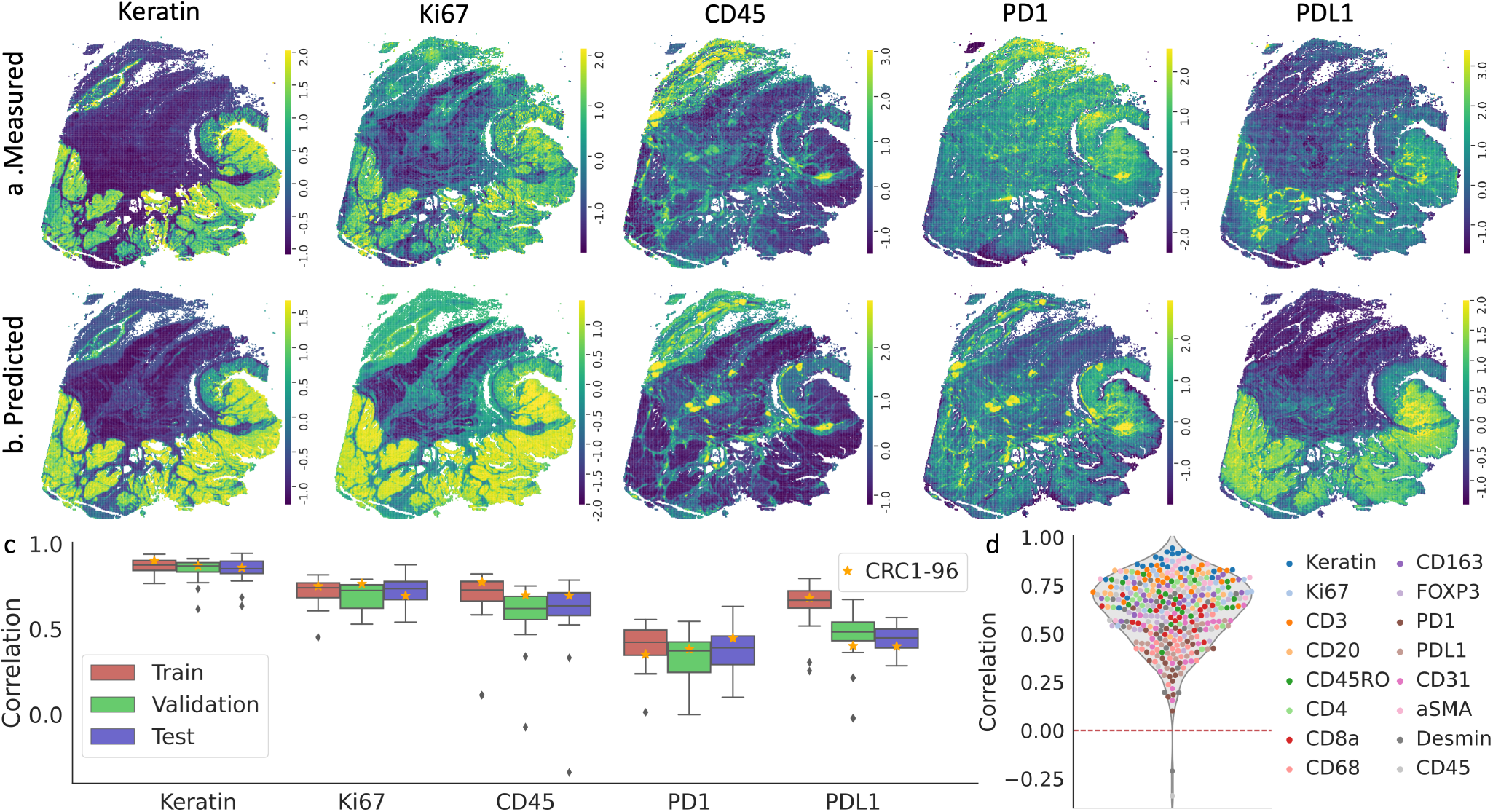
Selected results on sample CRC01 slice 96 for five markers: Keratin, Ki67, CD45, PD1, PDL1. (a) tile level CyCIF expression (z-score). (b) Prediction of our model for each tile (z-score).(c) Pearson correlation between measured and predicted values. We calculate the correlations for the train/validations/test tiles of each slice of sample CRC01 separately (slice 96 is marked).

On the other hand, some markers, such as PD1 or PDL1, are more difficult to correctly infer from the H&E image although the model’s predictions are still positively correlated to the measured expression. The median correlation for PD1 is 0.4 with no difference between train and test sets, while for PDL1 we see a slight degradation in correlation between train and test sets. The measured values of these markers have grid like artifacts as a result of the acquisition cycles and image stitching of the CyCIF protocol [8]. This phenomenon contributes to the lower correlation between the measured and predicted values. However, we still see that the model has good performance capturing highly expressed tiles (Figure S4, Table S1). The top-20% accuracy for PD1 and PDL1 in slice 96 was 38% and 50% respectively.

### 2.3 HIPI generalizes to new colorectal samples

Next, we evaluated HIPI on tiles from 7 additional CRC samples that were not used for training and originated from different patients spanning different histologic and molecular subtypes (named *CRC* 2, 3, 12, 13, 14, 15 and 17 in [25], Figure 1f). These samples were selected based on the physical proximity of their H&E and CyCIF slices. We preprocessed the new samples similar to the training samples (Section 2.1). This out of distribution test dataset from the 7 new tumors yielded 794k tiles.

We see that HIPI predictions are mostly positively correlated with the marker measurements (Figure 3 and S2). Naturally, the predictive performance is lower in these out of distribution samples in comparison to the slides of patient CRC01 (Figures S2 and S3). Similar to the training slides from CRC01, we see considerable differences in predictive performance between markers. For example, the predictions of Ki67 and CD45 are highly correlated with the true CyCIF values across the new samples achieving levels similar to the test sets of CRC01. HIPI correctly predicts the top-20% tiles with highest Ki67 expression with 46-77% accuracy, similar to CRC01 samples (Figure S4, Table S1). Whereas for CD45 we observe 38-57% highly expressed tile top-20% accuracy, slightly lower than CRC01 samples. Consistent with our observations on the training samples, PD1 and PDL1 have lower levels of overall correlations. However, these levels of correlations are similar to the range of correlations observed for CRC01 test sets (Figure S3). In addition, we see that HIPI predicts the highly expressed tiles with 29-44% and 20-50% accuracy for PD1 and PDL1 respectively (Figure S4, Table S1).

**Figure 3:**
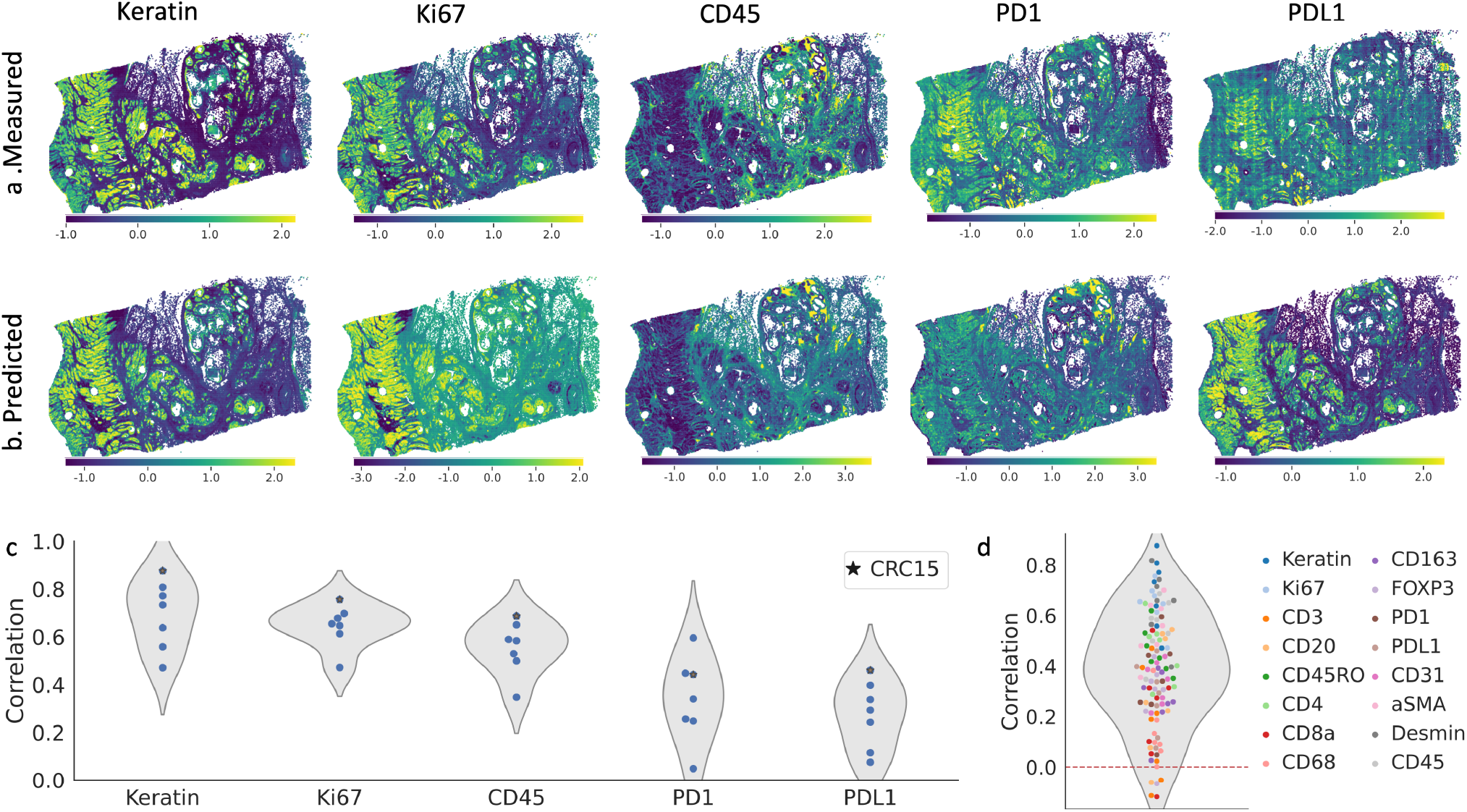
Selected results on sample CRC15 for five markers: Keratin, Ki67, CD45, PD1, PDL1. (a-b) similar to to 2(a-b). (c) Pearson correlation between measured and predicted values for all external CRC samples (sample CRC15 is marked).

### 2.4 HIPI Captures Cell Types and Marker Co-occurrence

Different cell types express different surface markers depending on the tissue morphology and other proximal cell types. The response of immune cells in the tumor boundary can have significantly influence disease progression and treatment response. Moreover, interactions between cells expressing certain markers such as PD1 and PDL1 can be therapeutically targeted and thus is clinically significant. It is thus important to determine the co-localization of different cell types and surface markers. Here, we analyze how tile level protein expression values capture cell surface marker expression and identify the co-localization of different cell types. Furthermore, we examine how HIPI predictions on H&E image tiles recapitulate the measured spatial relationship. We note that while our analysis is at the tile level and not in single cell resolution, it is sufficient to observe co-occurrence (e.g. PD1 and PDL1) of marker expression at the same tile to infer cell interaction.

We binarized the normalized protein expression values to obtain marker calls for each tile. To that end, we fit a two component Gaussian Mixture Model on the values of each protein and slide separately, similar to [25]. We see that the tile level marker calls capture the abundance and distribution of cell level CyCIF calls (Figure 4a in comparison to Figures 7abcdk of [25]). This shows that the tile level calls can also potentially recover the interaction or co-localization of different cell types within a tile.

**Figure 4:**
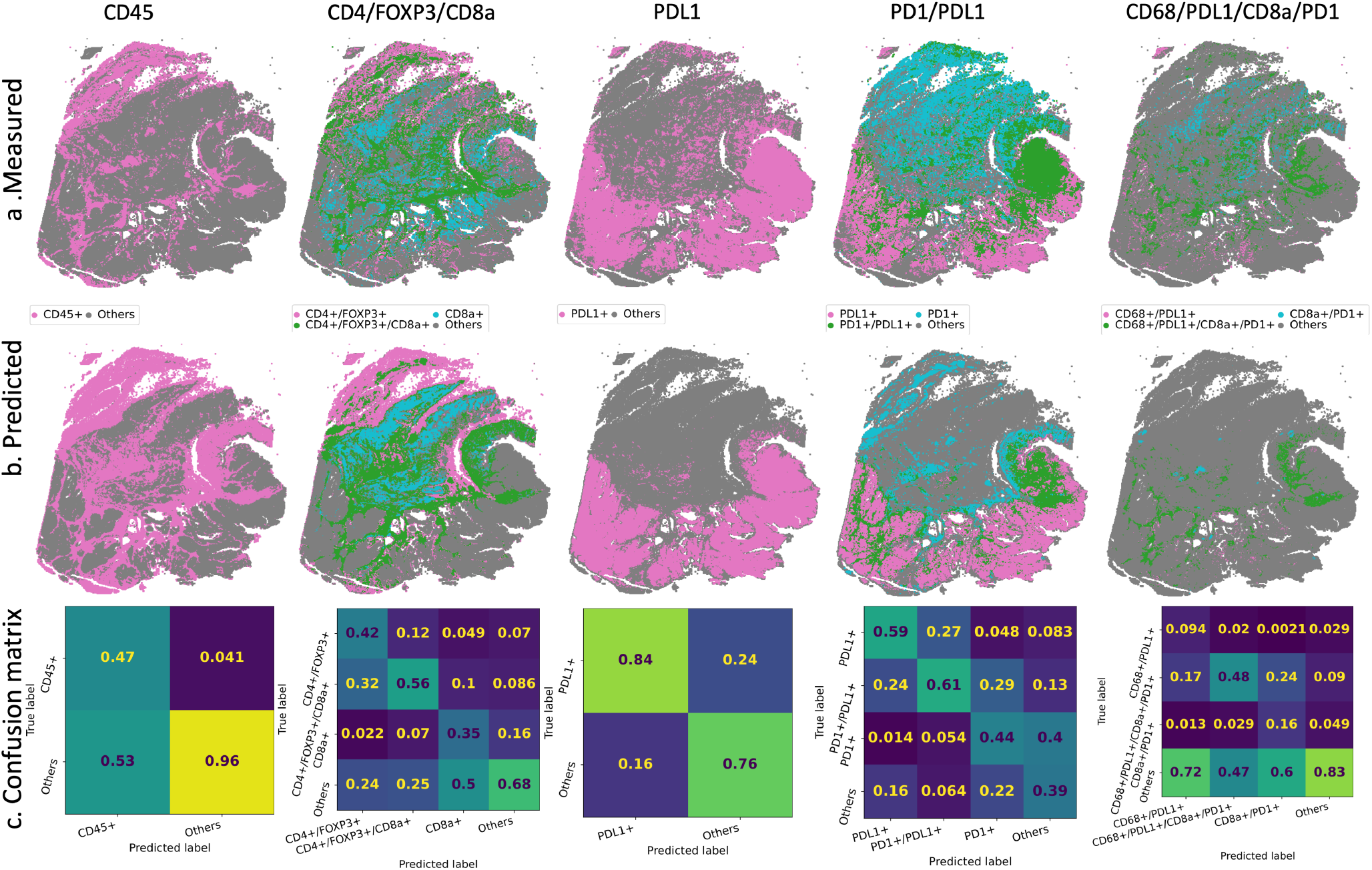
Selected cell marker occurrence in CRC01-96. (a) Occurrence and co-occurrence of CD45, CD4/FOXP3/CD8a, PDL1, PD1/PDL1 and CD68/PDL1/CD8a/PD1 in measured tile level data. (b) Predicted occurrence by HIPI. (c) Confusion matrices classifying different marker co-occurrence. Matrices are normalized column-wise by the total number of predictions for each class. Therefore, the inline numbers give the fraction of tiles predicted by the model to be in a certain class which truly belong to that class.

**Figure 5:**
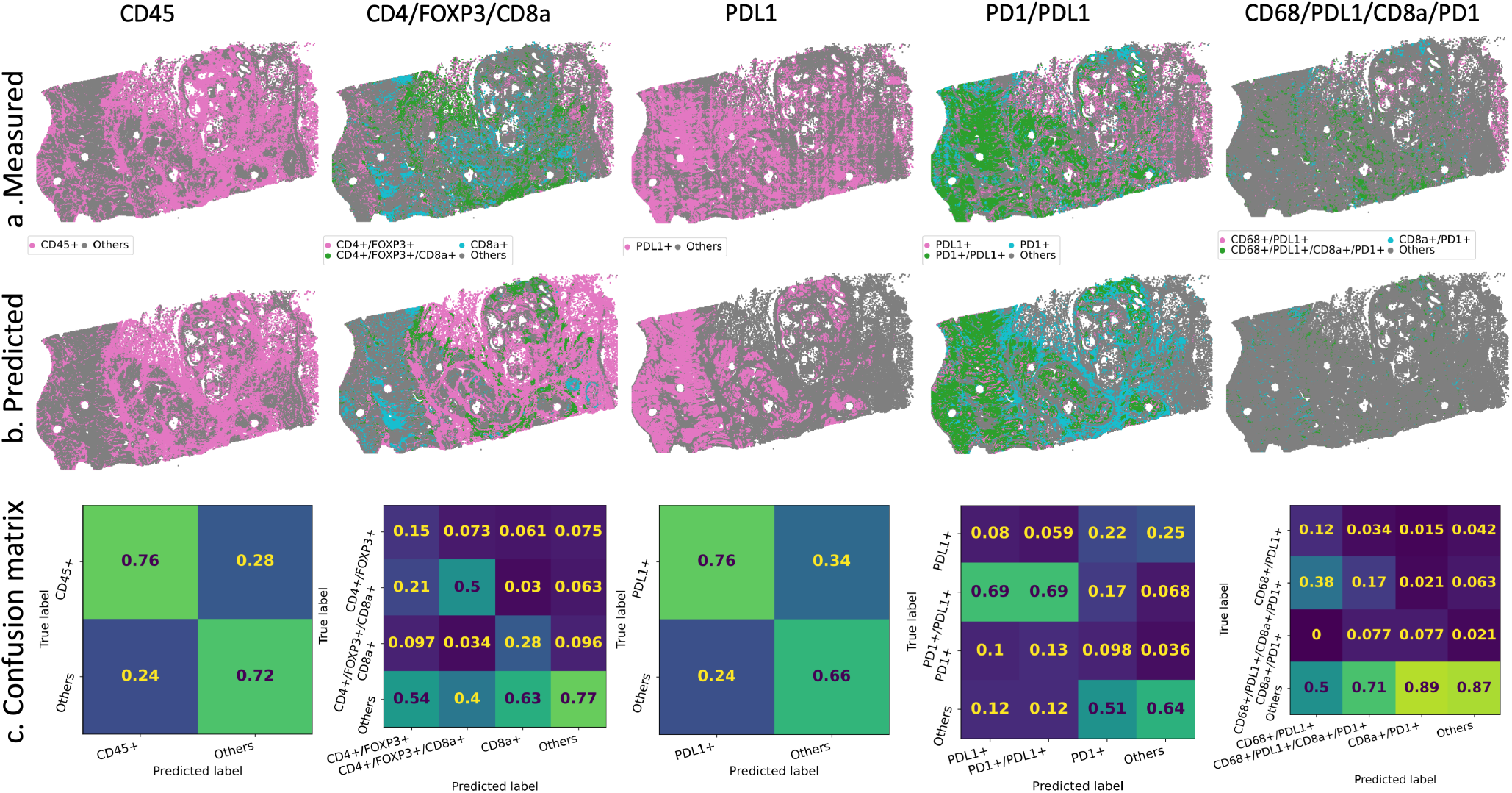
Cell markers occurrence in CRC15. See Figure 4 for panel description.

Next, we evaluated the performance of HIPI in calling marker distribution and co-occurrence (Figure 4b). We see that HIPI predictions provide smoother marker calls in comparison to the measured tile level calls. For instance, we do not see the grid-like artifacts observed in the measurements of some markers such as PDL1. The model tends to be conservative and give high precision in calling the true marker occurrences, where the precision is defined as the number of true positive calls divided by the total number of predicted positive calls. That is, tiles called by the model to be marker positive are likely to be true marker positive (Figure 4c). In particular, we see that marker co-occurrences are called with high precision. For example, the precision recovering CD4+/FOXP3+/CD8a+, PD1+/PDL1+ and CD68+/PDL1+/CD8a+/PD1+ markers is 0.56, 0.61 and 0.48 respectively, even better than the precision recovering the individual markers. We further observe that the model predictions underscore the immunosuppressive interaction between PD1+ and PDL1+. We see that tiles of PD1+:PDL1+ interactions are enriched for CD45+ (3.0 *∗* 10^*−*16^ one-sided paired t-test p-value) and depleted for CK+ (1.1 *∗* 10^*−*13^ one-sided paired t-test p-value) in comparison to non-interacting tiles (Figure S5). Moreover, the model highlights these trends even better than the tile level measurements for individual slices improving the statistical significance.

We further evaluated HIPI on marker prediction and co-occurrence on the external test CRC samples that were not used for training (Section 2.3). We see that the model is able to predict CD45 and PDL1 with 0.76 precision, removing unwanted artifacts in the measured data. Similarly, we are able to predict the PD1+/PDL1+ and CD4+/FOXP3+/CD8a+ interactions with 0.69 and 0.5 precision respectively. Since cell markers overlap, the model sometimes predicts a partial marker set. For example, out of the tiles HIPI predicted to be CD4+/FOXP3+, 21% were measured to be CD4+/FOXP3+/CD8a+. In addition, we see that HIPI captures the decrease of Keratin expression (0.03 one-sided paired t-test p-value) and the increase in CD45 expression (0.01 one-sided paired t-test p-value) in PD1+:PDL1+ interacting tiles in the new samples (Figure S5). In contrast, the measured tile level calls do not show similar trends which further illustrates the advantage of out model.

### 2.5 Comparison to

In addition to HIPI, we also trained *baseline* model uses only the mean image intensities together with a linear layer. That is, the baseline model does not use the tissue morphology but only the color intensities in tiles. We used the same training tiles from samples of CRC01 and evaluated the performance on the left out tiles and the tiles of the other CRC samples.

We observe that our model which uses a pretrained SSL feature extractor, consistently gives higher correlation to the measured tile protein expression than baseline model that uses only color intensities (Figure S1). Each model shows similar correlation metrics between tiles from the training, validation and test sets. Some markers are relatively easy to identify with relatively high correlation even with the model (Figure S1). For example, the baseline model gives a median correlation of 0.7 on Keratin levels whereas the HIPI model gives a median correlation of 0.86. On the other hand, there are proteins that are relatively difficult to predict based only on an H&E image tile. For instance, prediction of CD31 expression has median correlations using the baseline models and a 0.45 median correlation using the HIPI model. For some proteins the HIPI model shows a significant advantage. For example, the baseline models gives median correlations of predicting CD45 while the HIPI model gives a 0.65 median correlation (Figure S1).

To evaluate the ability to generalize to new samples, we evaluate the baseline models performance on the tiles of the additional CRC samples [25]. We see that our HIPI model is able to generalize to new samples correctly predicting tile level expression of several proteins (Figures S2 and S3). While some proteins can be predicted on the new samples at similar levels of correlation as the training samples (e.g. Keratin, Desmin Ki67, CD45), other proteins are more difficult to predict (e.g. CD3, CD20, CD68). Still, we see that our model is able to better predict tile protein expression than the baseline models. This shows that the image tissue structure indeed provides valuable information (Figures S2 and S3).

## 3 Discussion

We describe HIPI, a method for predicting protein markers measured with CyCIF from H&E images. We specifically described our results on colorectal cancer samples. We design a pipeline for aligning and processing paired CyCIF and H&E images of adjacent tumor slices. We then train a model, based on a dedicated pre-trained H&E image encoder, for predicting CyCIF protein expression from small image tiles. We evaluate our model on unseen image regions from samples used for training as well as new samples never used in training. This is in contrast to previous methods for generating spatial molecular measurements from H&Es, which only evaluate on samples or regions coming from the same samples or regions used for training their models; as a result, much less can be said about such methods’ generalization abilities. On most measured proteins, our model is able to recover the expression levels on both internal and external evaluation samples. This shows that tissue morphology captured in the H&E image provides important information pertaining to the types of cells observed, the markers they express, and the extent of this expression. AI inference of protein expression from H&E can be used as a first stage assessment and that actual molecular assay of IHC staining can follow to obtain high confidence for the suggested findings.

Unlike CyCIF or other technologies, HIPI predicts marker expression at low resolution image tile level. This lower resolution tile level measurement is an aggregate of all cells in the frame. As a result, the expression of different cells may cancel one another and may hinder the ability to observe rare cell types. In addition, with tile level view we lose information that might be important in order to understand cell-cell interactions. On the other hand, using tile level measurements allows our method to cope with small misalignments between the adjacent tissue slices used for training. Moreover, tile level measurements reduce the variation observed in cell level data or the exact calls of cell positions.

Our method can be improved and generalized in several ways. First, while we used CyCIF, there are other spatial technologies that yield molecular measurements at different resolutions [27]. Although methods have been proposed to generate spatial transcriptomics measurements from H&E images, they do not generalize to out of distribution samples [17]. Second, although we used our model to train and infer on colorectal cancer samples, we could train and generalize to other cancer types. In fact, the cancer type, as well as other clinical information can be incorporated into the model to discern inter-sample variability. Third, it might be useful to improve our spatial resolution to allow single cell resolution instead of tile level resolution. This requires better cell segmentation from both the H&E and the CyCIF images and calling the cell level CyCIF protein expressions. Moreover, it requires aligning not only the images but also the cells between two (slightly) different tissue slices.

## Data availability

Raw H&E images, raw CyCIF images and processed CyCIF cell data can be obtained from https://github.com/labsyspharm/CRC_atlas_2022.

## 4 Author Contribution

R.Z. developed the method and performed the analyses. R.Z., L.A., Z.Y. and D.F. conceived and designed the method and analyses. All authors read and approved the final manuscript.

## S1 Supplementary Methods

### S1.1 Preprocessing CyCIF Samples

#### S1.1.1 Aligning H&E and CyCIF

The same tissue section cannot be used to produce both histological stained image and immunofluorescence image. Therefore, usually multiple thin serial sections of the same tissue are generated and subjected to staining or immunofluorescence imaging. However, adjacent slices and their corresponding images are not perfect replicates of one another. First, there are some physical differences between sections caused by natural tissue structure as well as deformations caused by the manual resection. Second, positioning of the slices for scanning and artifacts created by the imaging technology result in unaligned images of adjacent sections.

We employ a heuristic alogirthm to align H&E and CyCIF images taken from consecutive tissue sections. Since the placement and orientation of the tissue varies between sections, we first find a global registration of the two images. To accomplish this, we apply a linear affine alignment on down-sampled grayscale versions of the H&E and CyCIF images. Because serial sections of the tissue are not exact copies of one another, we then refine our registration on small local tiles. We align overlapping local tiles using a non-linear registration technique to account for small tissue differences.

Let 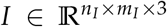 and 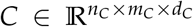be the H&E and CyCIF images respectfully. In addition, we assume the CyCIF data was processed to produce cell-level expression values. Let *W* ∈ *ℕ*^*l×*2^ be the coordinates of *l* cells in the CyCIF image (i.e. for each cell *i, W*[*i*] ∈ [0 … *n*_*C*_] *×* [0 … *m*_*C*_]) and let 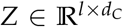 be the expression values for each cell. Our goal is to find translated coordinates *W* ′ ∈ *ℕ*^*l×*2^ of the cell, where *W ′* [*i*] ∈ [0 … *n*_*I*_] *×* [0 … *m*_*I*_] is now the approximated position of the cell in the H&E image.

We apply the following alignment heuristic (Figure 1b): First, we grayscale the H&E image. Then, we resize the CyCIF image to around the H&E scale and use one of the auto autofluorescence channels to grayscale the CyCIF image. We find a global affine alignment transformation on downsampled versions of the two images and translate the cell coordinates accordingly. Finally, we perform local non-linear alignment of images tiles using both affine and B-spline registration. We transform the coordinates of the cells to the H&E image by finding an approximate inverse transformation.

#### S1.1.2 Extracting tile Level Expression

Let *W* ′ ∈ *ℕ*^*l×*2^ be the coordinates of *l* cells in the H&E image and let *Z* 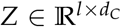 be the expression values for each cell. We log transform the expression values and normalize each protein by its mean and standard deviation to obtain *Zℕ* (Figure 1c). We extract strided image tiles of size *w* and the corresponding cells in each tile. For each such tile with a predefined minimum number of cells, we calculate the mean expression for each protein of cells within a tile (Figure 1c). We denote by 𝒫= (*I*_*i*_, *z*_*i*_), the set of H&E image tiles and corresponding expression vectors pairs.

### S1.2 Model for Predicting Expression from H&E tiles

Letℱ: [0, 1]^*w×w×*3^ *→ ℝ*^*d*^ be a function that transforms a *w × w* H&E image to a *d*-dimensional feature vector and let 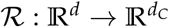 be a function that maps a *d*-dimensional feature vector to a normalized tile expression of *d*_*C*_ proteins. Our goal is to train a model 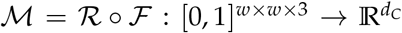 that predicts a normalized tile expression of *d*_*C*_ proteins from a *w × w* H&E image and minimizes the squared error 𝔼_*I,z∈P*_ (*1M*(*I*) *− z1*).

Self-Supervised learning (SSL) methods, which leverage large amounts of unlabeled data to pretrain meaningful feature extractors, have been shown to out-performs supervised pre-training on multiple task [21]. This approach is especially compelling in the field of digital pathology where there are large data sets of unlabeled slides in comparison to limited number of annotated data sets. Indeed, SSL methods for digital pathology have been shown as meaningful feature extractors for multiple downstream tasks even in the face of limited amount of labeled data [22, 23].

Here we use an SSL model pre-trained on publicly available pathology images and use it as a fixed feature extractor ℱ [23]. Specifically, we use a Vision Transformer (ViT) trained using DINO which gave good results on various task in a recent benchmark [28, 21]. On top of the feature extractor, we train a regressor ℛmodeled as a fully connected multi-layer perceptron. We call our combined model HIPI.

For reference, we also train baseline model extracts the mean color intensities and uses a linear layer to predict the expression.

### S1.3 Implementation Details

#### Global alignment of H&E and CyCIF

We resize the CyCIF image by 2 to grossly match the H&E image resolution and use the first *Hoechst* channel for grayscaling. We find global affine registration on *×*0.01 downsized images using Elastix [29].

#### Local alignment of H&E and CyCIF

We go over the images with tiles of size 2^14^ pixels and a 2^10^ stride while allowing of 2^9^ extra pixels slack in the CyCif image. We only aligned tiles that had at least 500 cells in them. To align tiles, we first downsize the tile by *×*2^*−*5^, the find an affine registration followed by a non-linear BSpline registration implemented with SimpleITK [29]. Since the BSpline transformation is not necessarily invertable, in order to translate the cell coordinate positions in the aligned images, we iteratively search for the closest inverse coordinate for each cell. Finally, since each cell may appear in multiple tiles, we take the final cell coordinates as the mean over all tiles the cell appears in.

#### Extracting tile level expression

We extract image tiles of size *w* = 256 pixels with a stride of 128 together with the corresponding cells in each tile. An image tile of size 256 pixels corresponds to roughly a 128*µm* tissue tile. We keep only tiles that contain at least 1 cells.

#### Pre-trained feature extractor

We use a pre-trained ViT that uses an internal tile size of 8 pixels and has 21.7M parameters [23]. We resize image tiles to 224 pixels to match the model’s design. The model outputs a feature vector of size *d* = 384.

#### Expression regressor

We use a multi-layer perceptron with 128, 64, 32, 16 hidden layers, a batch norm layer, a GELU activation and a 0.2 dropout layer. The model has 60.9K trainable parameters.

#### Training

We use a base learning rate of 10^*−*6^, batch size of 512 and an Adam optimizer with default parameters. We randomly augment the image tiles during training to allow for better generalization using a recent augmentation scheme *RandStainNA* designed specifically for pathology images [26]. Furthermore, we randomly flip and rotate image tiles in contrast to the common practice on natural images.

## S2 Supplementary Analysis

**Figure S1:**
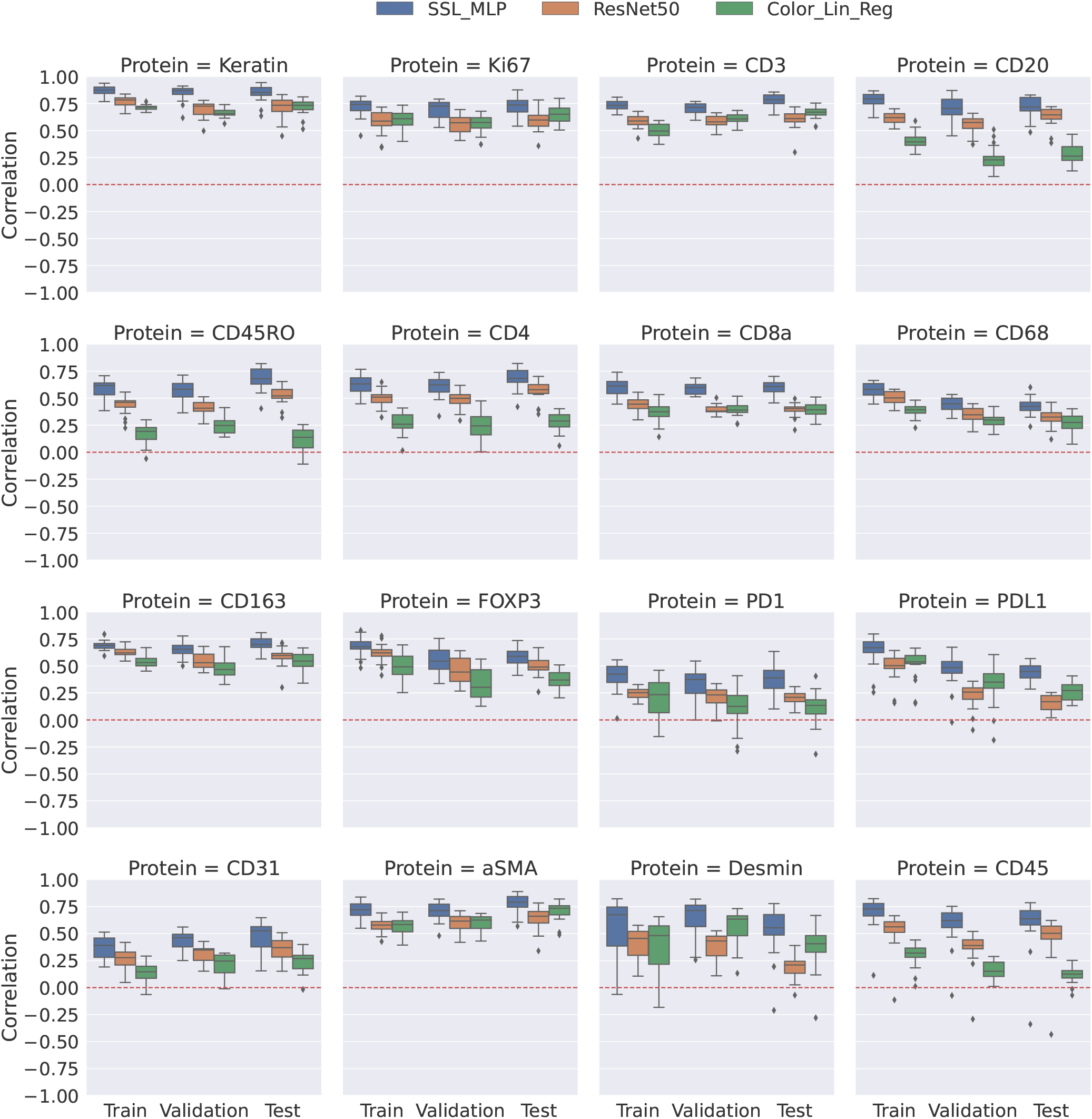
Correlation between measured and predicted tile level expression for 22 samples from patient CRC01 [25]. Results are divided between tiles used for training and validating models, and tiles left out for testing. Correlations for HIPI are shown in blue, correlations for the baseline are shown in orange

**Figure S2:**
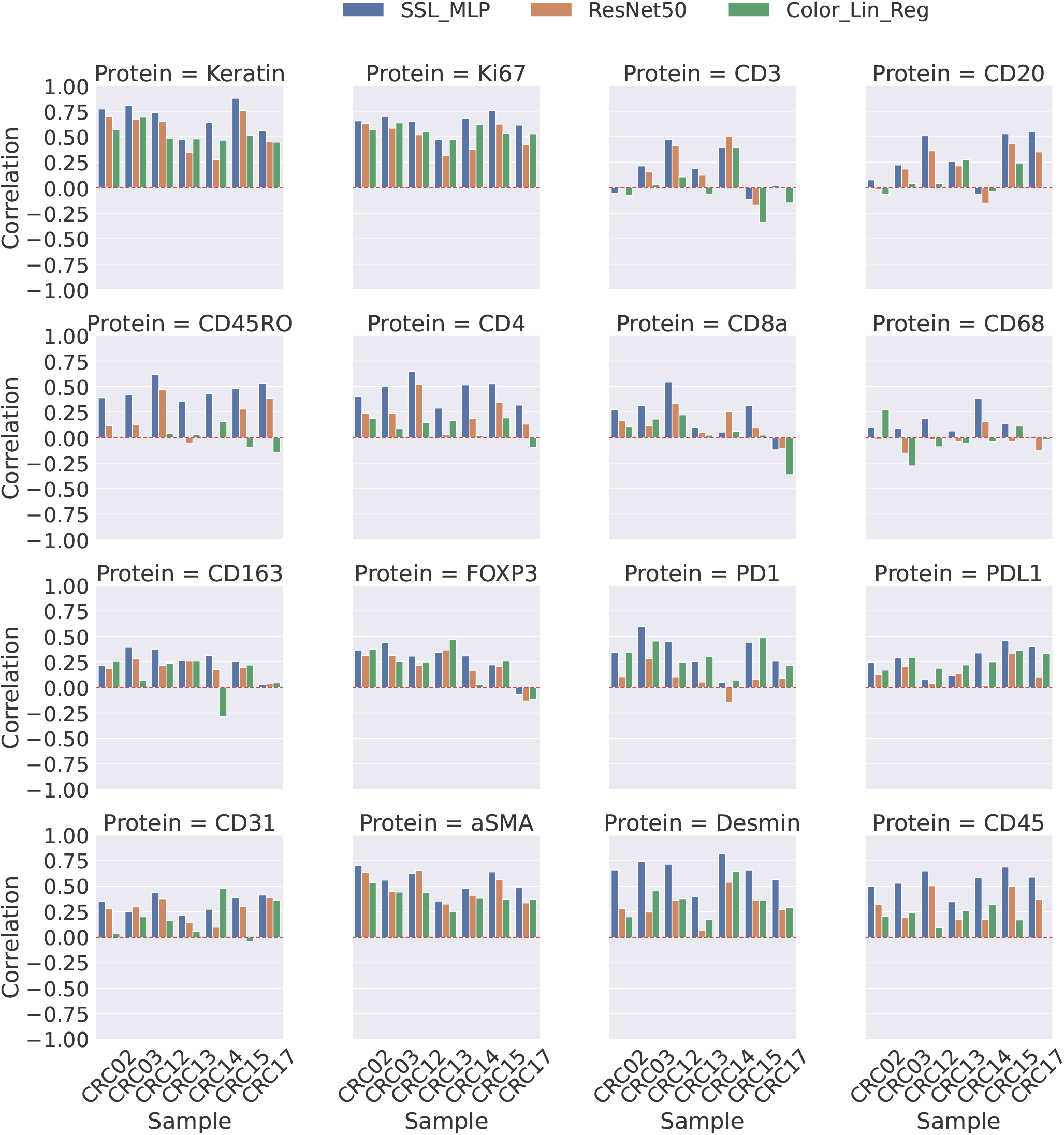
Correlation between measured and predicted tile level expression for patients CRC 2, 3, 12, 13, 14, 15 and 17. [25].

**Figure S3:**
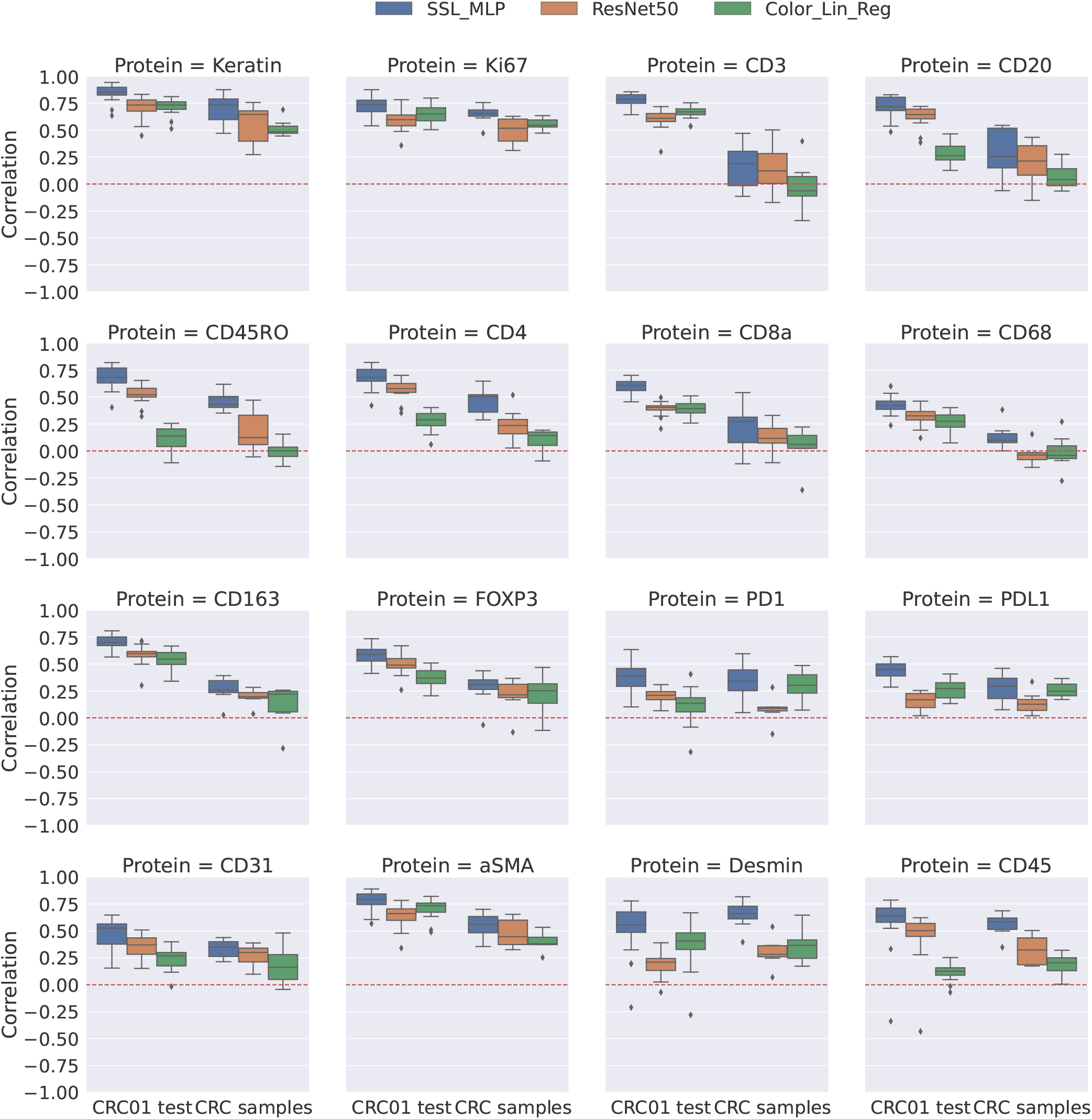
Correlation between measured and predicted tile level expression for test tiles of CRC01 and patients CRC 2, 3, 12, 13, 14, 15 and 17. [25].

**Figure S4:**
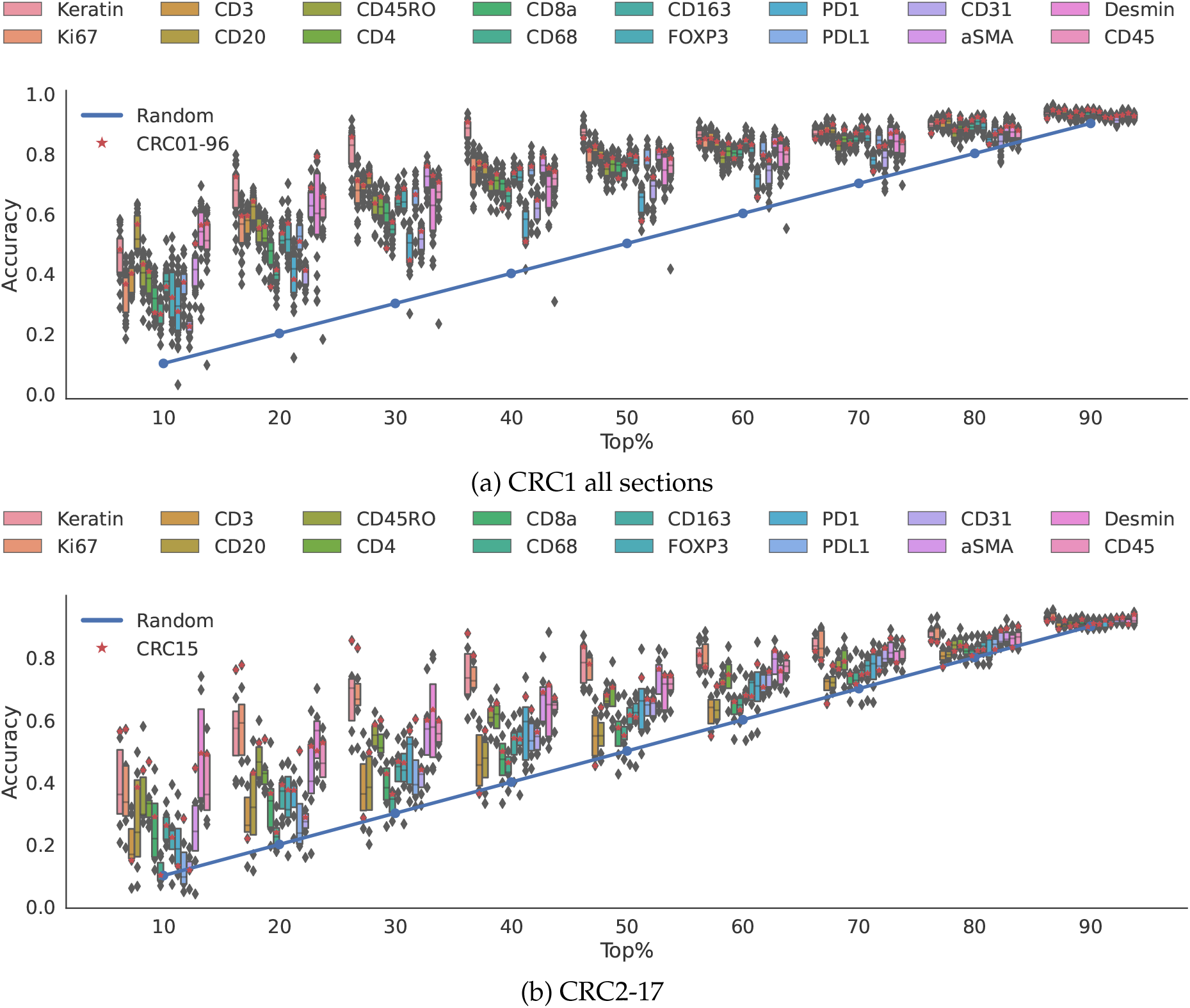
HIPI top X% accuracy. For each slide and marker, we calculate the overlap between the top X% of measured tiles and the top X% of predicted tiles and normalize by X%. (a) Slides form patient CRC1 where slide 96 is marked in red. (b) Slides from patients CRC2-17 where the slide from CRC15 is marked in red. Blue line represents the random chance accuracy.

**Table S1:** Top-20% accuracy. Accuracy % of HIPI in predicting the 20% of tiles with the highest expression for each sample and protein.

| Sample | Keratin | Ki67 | CD3 | CD20 | CD45RO | CD4 | CD8a | CD68 | CD163 | FOXP3 | PD1 | PDL1 | CD31 | aSMA | Desmin | CD45 |
| --- | --- | --- | --- | --- | --- | --- | --- | --- | --- | --- | --- | --- | --- | --- | --- | --- |
| CRC01-1 | 60 | 50 | 51 | 59 | 50 | 46 | 53 | 31 | 48 | 46 | 43 | 52 | 40 | 59 | 55 | 58 |
| CRC01-6 | 47 | 43 | 49 | 55 | 41 | 37 | 43 | 29 | 52 | 56 | 32 | 48 | 31 | 53 | 56 | 52 |
| CRC01-13 | 51 | 46 | 53 | 57 | 48 | 50 | 49 | 36 | 52 | 45 | 32 | 43 | 33 | 59 | 39 | 54 |
| CRC01-19 | 56 | 36 | 53 | 56 | 50 | 50 | 49 | 39 | 48 | 49 | 32 | 46 | 36 | 62 | 30 | 54 |
| CRC01-24 | 59 | 50 | 56 | 66 | 52 | 50 | 45 | 37 | 50 | 52 | 29 | 43 | 37 | 58 | 34 | 60 |
| CRC01-28 | 64 | 43 | 52 | 62 | 53 | 51 | 38 | 39 | 56 | 52 | 35 | 36 | 36 | 60 | 69 | 59 |
| CRC01-33 | 76 | 58 | 58 | 62 | 59 | 52 | 42 | 40 | 55 | 64 | 42 | 57 | 41 | 65 | 73 | 64 |
| CRC01-38 | 66 | 56 | 57 | 63 | 55 | 51 | 49 | 39 | 50 | 54 | 40 | 52 | 38 | 67 | 56 | 63 |
| CRC01-43 | 73 | 50 | 61 | 67 | 61 | 60 | 48 | 45 | 63 | 60 | 46 | 42 | 49 | 74 | 53 | 65 |
| CRC01-48 | 71 | 55 | 61 | 66 | 58 | 57 | 47 | 43 | 49 | 46 | 50 | 53 | 42 | 66 | 58 | 66 |
| CRC01-53 | 61 | 56 | 51 | 57 | 50 | 45 | 46 | 35 | 51 | 50 | 27 | 49 | 36 | 54 | 45 | 59 |
| CRC01-58 | 76 | 65 | 61 | 65 | 57 | 58 | 50 | 39 | 50 | 57 | 44 | 58 | 40 | 65 | 30 | 68 |
| CRC01-63 | 76 | 60 | 63 | 65 | 58 | 52 | 56 | 43 | 51 | 69 | 11 | 59 | 42 | 64 | 61 | 18 |
| CRC01-68 | 77 | 59 | 57 | 65 | 59 | 59 | 53 | 38 | 53 | 50 | 40 | 53 | 43 | 67 | 72 | 68 |
| CRC01-73 | 61 | 56 | 51 | 55 | 41 | 44 | 36 | 40 | 55 | 43 | 35 | 55 | 34 | 54 | 73 | 51 |
| CRC01-77 | 66 | 62 | 57 | 62 | 49 | 50 | 37 | 42 | 49 | 41 | 42 | 51 | 42 | 58 | 68 | 59 |
| CRC01-83 | 68 | 51 | 57 | 57 | 51 | 50 | 38 | 39 | 47 | 47 | 46 | 56 | 41 | 60 | 77 | 61 |
| CRC01-85 | 65 | 39 | 58 | 57 | 50 | 49 | 51 | 42 | 48 | 56 | 43 | 59 | 35 | 58 | 73 | 61 |
| CRC01-90 | 79 | 66 | 60 | 68 | 60 | 55 | 57 | 40 | 57 | 59 | 48 | 59 | 41 | 69 | 52 | 67 |
| CRC01-96 | 72 | 59 | 59 | 64 | 55 | 55 | 35 | 41 | 53 | 56 | 38 | 50 | 41 | 68 | 78 | 65 |
| CRC01-101 | 70 | 64 | 58 | 62 | 55 | 53 | 47 | 34 | 48 | 47 | 50 | 54 | 36 | 62 | 77 | 63 |
| CRC01-105 | 72 | 58 | 60 | 62 | 55 | 53 | 47 | 45 | 49 | 45 | 45 | 48 | 42 | 67 | 79 | 65 |
| CRC02 | 56 | 46 | 12 | 16 | 41 | 37 | 33 | 18 | 26 | 30 | 39 | 20 | 26 | 62 | 45 | 38 |
| CRC03 | 58 | 59 | 31 | 29 | 41 | 42 | 39 | 20 | 45 | 45 | 38 | 23 | 24 | 40 | 70 | 46 |
| CRC12 | 67 | 70 | 38 | 41 | 59 | 44 | 52 | 22 | 34 | 27 | 44 | 19 | 35 | 38 | 58 | 57 |
| CRC13 | 40 | 40 | 26 | 31 | 39 | 36 | 25 | 26 | 37 | 46 | 21 | 20 | 27 | 34 | 40 | 39 |
| CRC14 | 57 | 58 | 46 | 11 | 46 | 40 | 25 | 32 | 40 | 31 | 29 | 40 | 15 | 17 | 60 | 43 |
| CRC15 | 76 | 77 | 21 | 42 | 52 | 53 | 36 | 23 | 39 | 37 | 37 | 50 | 28 | 51 | 50 | 52 |
| CRC17 | 39 | 50 | 26 | 53 | 49 | 42 | 19 | 17 | 28 | 16 | 34 | 25 | 30 | 52 | 56 | 53 |

**Figure S5:**
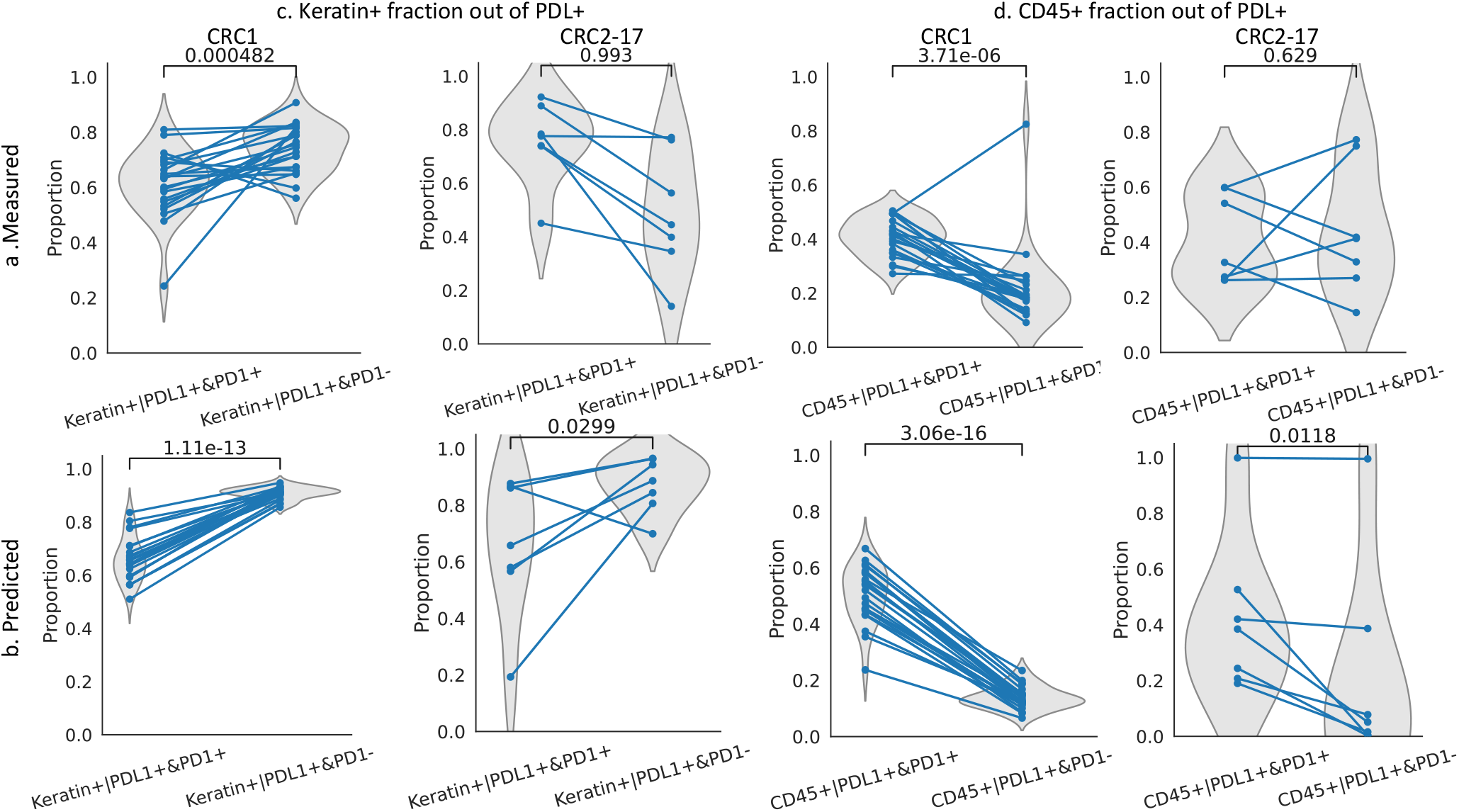
The proportion of Keratin+ and CD45+ tiles out of PDL1+:PD1+ interacting and non-interacting tiles. (a) Proportions observed in the processed measured tiles. (b) Proportions observed the inferred tiles. Results are summarized for each slice and presented for the slices of sample CRC1 and and slices coming from the other patients CRC2-17. The p-value of the one-sided paired t-test for the increase/decrease between PDL1+:PD1+ interacting and non-interacting tiles is presented above.

